# Memory and Hippocampal Responses to Event Boundaries are Modulated by Global Brain States

**DOI:** 10.64898/2026.04.14.718575

**Authors:** Dora Gozukara, Nasir Ahmad, Djamari Oetringer, Linda Geerligs

## Abstract

Our daily experiences unfold as a continuous stream, yet we perceive and remember them as discrete events. Event boundaries, the moments of transition between these events, are known to elicit increases in hippocampal activity believed to reflect memory encoding. However, it remains unknown how this hippocampal response relates to large-scale brain dynamics. Here, using fMRI data from two independent datasets (Sherlock and StudyForrest), we applied the Greedy State Boundary Search (GSBS) algorithm to whole-brain activity patterns and identified two recurring global brain states corresponding to the Default Mode Network (DMN) and Task-Positive Network (TPN). We found that event boundaries were associated with an increased probability of being in the TPN state, and that hippocampal activity was generally higher during TPN states. The hippocampal response to event boundaries appeared predominantly during TPN states. When overall state-related differences in baseline hippocampal activity were controlled for, event boundaries elicited a hippocampal response regardless of the concurrent global state. Critically, individual differences in the tendency to shift toward the TPN state at event boundaries; as well as overall time spent at the DMN state predicted subsequent memory for narrative content, whereas univariate hippocampal activity at boundaries did not. These findings demonstrate that hippocampal event boundary responses are modulated by global brain state dynamics, and suggest that the interplay between large-scale network configurations and event segmentation plays a key role in how continuous experience is encoded into memory.

## 1 Introduction

Our daily experiences unfold as a continuous stream of sensory information, yet we seem to perceive and remember them as discrete events. This fundamental aspect of human cognition, termed event segmentation, organises ongoing experience into meaningful units that can be understood, stored, and later retrieved (Kurby & Zacks, 2008; Zacks & Swallow, 2007). According to Event Segmentation Theory (EST), individuals continuously maintain mental representations of the current situation that are updated when significant changes occur in the environment, such as shifts in location, characters, goals, or temporal context (Radvansky & Zacks, 2017; Zacks & Swallow, 2007). These moments of change, known as event boundaries, have disproportionate effects on how we perceive and remember our experiences.

It is well established that the hippocampus plays a critical role in processing event boundaries during continuous experience. Research has shown that hippocampal activity reliably increases at event boundaries during movie watching, with this response being both sensitive and specific to boundary moments (Ben-Yakov & Dudai, 2011; Ben-Yakov & Henson, 2018; Ben-Yakov et al., 2013). Specifically, hippocampal activity following an event predicts the strength of neural state reactivation during subsequent recall (Baldassano et al., 2017), and this response at boundaries is modulated by boundary salience (Ben-Yakov & Henson, 2018). This hippocampal engagement at event boundaries has been proposed to reflect the binding of event representations into episodic memory traces, effectively serving as the brain’s “film editor” which structures continuous experience into memorable episodes (Ben-Yakov & Dudai, 2011; Ben-Yakov & Henson, 2018; Silva et al., 2019).

To investigate the neural underpinnings of event segmentation and its role on memory, recent research has focused on the neural state boundaries which tend to accompany event boundaries (Baldassano et al., 2017; Geerligs et al., 2022; Oetringer et al., 2025). Neural states are defined as periods of relatively stable neural activity patterns that follow one another with rapid pattern changes in between (Baldassano et al., 2017; Geerligs et al., 2022). These states can be identified at multiple spatial scales. At the local level, neural states in individual brain areas have been shown to form a nested temporal hierarchy; where lower-level perceptual features give rise to shorter and faster states in primary sensory areas, while higher-level semantic and scene-level features give rise to longer states in higher-order associative areas (Baldassano et al., 2017; Geerligs et al., 2022; Gözükara et al., 2026; Oetringer et al., 2024). At the global level, whole-brain activity patterns show similar discrete transitions which reflect coordinated shifts across distributed functional networks (Gözükara et al., 2026; Thompson et al., 2014; Yamashita et al., 2021). Our recent work has shown that local and global neural states are related by dissociable, and that event segmentation is linked to neural state transitions at both scales (Gözükara et al., 2026). While it has been established in previous research that local neural state boundaries are associated with hippocampal response and memory encoding (Baldassano et al., 2017; Ben-Yakov & Henson, 2018), the role of global brain states in these processes has not been investigated.

To understand what these global brain states reflect, it is important to consider the brain’s large-scale functional organisation. On the most coarse level of organization, the human brain is intrinsically organised into anti-correlated networks: the Default Mode Network (DMN) and task-positive networks (TPN), sometimes referred to as the dorsal attention network or central executive network (Fox et al., 2005; Raichle et al., 2001). The DMN, comprising regions such as the medial prefrontal cortex, posterior cingulate cortex, and angular gyrus, is typically active during internally-directed cognition, including autobiographical memory retrieval, future thinking, and social cognition (Buckner et al., 2008; Menon, 2023; Spreng et al., 2009). Conversely, the TPN, encompassing frontoparietal regions, is engaged during externally-directed attention and cognitive control (Fox et al., 2005). These two networks have been shown to exhibit an anti-correlated relationship, such that activation of one network is accompanied by suppression of the other (Gözükara et al., 2026; Thompson et al., 2014). During rest and naturalistic stimulation, global neural states have been shown to correspond to these two competing network configurations, with whole-brain activity alternating between DMN-dominant and TPN-dominant states, potentially supporting the brain’s ability to flexibly shift between internal and external processing modes (Gözükara et al., 2026).

This large scale organisation leads to questions regarding the hippocampal response at event boundaries. On one hand, event boundaries are associated with increased hippocampal activity (Ben-Yakov & Henson, 2018) and are thought to trigger attentional orienting responses (Zacks & Swallow, 2007), which would predict a shift toward TPN dominant processing. On the other hand, the hippocampus shows strong functional connectivity with DMN regions and is often considered part of the extended DMN (Ranganath & Ritchey, 2012; Rugg & Vilberg, 2013), while the DMN itself has been proposed to support memory consolidation, narrative integration, and the construction of coherent internal models (Kaefer et al., 2022; Menon, 2023). If event boundaries simultaneously engage attentional networks and trigger hippocampal memory encoding, it is unclear whether the hippocampal boundary responses, as well as accompanying memory processes, are associated with DMN-dominant or TPN-dominant global states.

Despite growing interest in both event segmentation and large-scale brain dynamics, the relationship between global brain states and hippocampal memory processes during naturalistic perception has not been directly investigated. Previous work have linked local neural state boundaries to hippocampal responses and subsequent memory (Baldassano et al., 2017; Ben-Yakov & Henson, 2018; Reagh et al., 2020), has showed that global states during naturalistic stimuli correspond to DMN and TPN configurations (Gözükara et al., 2026), and that EEG derived whole-brain states relate to memory performance (Henderson et al., 2025). However whether the hippocampal response at event boundaries depends on the concurrent global brain state and whether global states modulate the link between event processing and memory remains an open question.

In the present study, we address this gap using fMRI data from two independent datasets: the Sherlock dataset (Chen et al., 2017) and the StudyForrest dataset (Hanke et al., 2016). We apply the GSBS algorithm to identify neural state transitions in whole-brain activity patterns and use density-based clustering to characterize recurring global brain states. Our analyses replicate the finding of two dominant global states corresponding to the DMN and TPN during naturalistic stimulation. We then examine whether the hippocampal response at event boundaries is modulated by the concurrent global brain state, testing if memory related hippocampal engagement preferentially occurs within one network configuration over the other.

We show that **(1)** event boundaries are associated with increased probability of being in the TPN state, potentially reflecting heightened external attention at moments of narrative change; **(2)** hippocampal activity is elevated at event boundaries, consistent with prior literature; and **(3)** the hippocampal response to event boundaries is modulated by the global brain state, with potentially larger responses when the brain is in a state conducive to memory encoding. Finally, we show that individual differences in global brain state dynamics and their relationship to event boundaries predict subsequent memory for the narrative content.

## 2 Methods

### 2.1 Datasets

To investigate how hippocampal activity at event boundaries is modulated by large-scale brain dynamics, we used two distinct datasets. We will describe each of these in turn as well as the preprocessing steps applied to each of them.

#### 2.1.1 StudyForrest dataset and preprocessing

The first dataset we used was the open access StudyForrest fMRI dataset. In this dataset 15 participants watched a 2-hour movie Forrest Gump dubbed in German (age 19–30, mean 22.4, 10 female, 5 male) (Hanke et al., 2016). The movie was split up into 8 parts and presented in two sessions on one day. This resulted in 3599 volumes of data acquired with a 32-channel head coil using a whole-body 3 Tesla Philips Achieva dStream MRI scanner. The images were T2*-weighted echo-planar images (gradient-echo, 2 s repetition time (TR), 30 ms echo time, 90° flip angle, 1943 Hz/Px bandwidth, parallel acquisition with sensitivity encoding (SENSE) reduction factor 2). There were 35 axial slices (thickness 3.0 mm) with 80 × 80 voxels (3.0 × 3.0 mm) of inplane resolution, 240 mm field-of-view (FoV), and an anterior-to-posterior phase encoding direction with a 10% inter-slice gap, recorded in ascending order. A T1-weighted image was acquired for anatomical alignment (Hanke et al., 2014).

The data was pre-processed as described in our previous work (Gözükara et al., 2026). Briefly, we started with data that was partially preprocessed by Liu et al. (2019). Their preprocessing included motion correction, slice timing, extracting the brain, applying a high-pass temporal filter of 1 Hz and denoising the data with ICA. We additionally performed alignment across runs, coregistration to the anatomical image and normalization to MNI space using SPM12 (Wellcome Centre for Human Neuroimaging, 2014). Per voxel and per run the data were z-scored. Data were masked to include only voxels active across all runs.

#### 2.1.2 Sherlock dataset and preprocessing

To be able to relate global activity patterns and hippocampal responses to memory performance, we also used the publicly available Sherlock dataset (Chen et al., 2017). In this dataset, 17 participants watched the first 48 minutes of Episode 1 of the BBC series Sherlock. The movie was split into two parts of roughly equal length (946 and 1030 TRs). During movie-watching, fMRI data were collected on a 3-T full-body scanner (Siemens Skyra) with a 20-channel head coil. Functional images were acquired using a T2*-weighted echo-planar imaging (EPI) pulse sequence (TR 1,500 ms, TE 28 ms, flip angle 64, whole-brain coverage 27 slices of 4 mm thickness, in-plane resolution 3 × 3 mm2, FOV 192 × 192 mm2), with ascending interleaved acquisition. Anatomical images were acquired using a T1-weighted MPRAGE pulse sequence (0.89 mm3 resolution).

We used the data as it was originally pre-processed by the authors (Chen et al., 2017). This included correcting for slice timing and head motion, high-pass filtering (140 s cutoff), coregistration to the anatomical image, transformation to MNI space and 6-mm smoothing. Afterwards, data were zscored across time at every voxel.

#### 2.1.3 ROI data

To be able to identify global brain states, we first computed the average timeseries for each ROI in the Schaefer 400 Atlas (Thomas Yeo et al., 2011) for each run and each participant in both datasets. The hippocampal signal was created by extracting data from the WFU PickAtlas toolbox (Maldjian et al., 2003). The hippocampal mask was resampled to match the resolution and dimensions of each participant’s MNI-space data prior to use as a region of interest. Data from each run was z-scored before concatenating it across runs for each participant.

#### 2.1.4 Behavioural data

For the StudyForrest dataset, event boundaries were defined by Ben-Yakov and Henson (2018), who had independent raters determine changes in abstract narrative events. There were 162 event boundaries in the movie. For the Sherlock dataset, 50 event (or scene) boundaries were identified as major shifts in the narrative (for example, location, topic, and/or time). For both datasets, a 4.5 second shift was applied to account for the hemodynamic lag between the neural and the behavioural data.

Participants recalled the Sherlock movie in as much detail as they could directly after watching it (Chen et al, 2017). They were instructed to recall events in the same order as the episode and to be as complete as possible. An event (or scene) was later rated as recalled if the participant described any part of the event. In our analyses we used the total number of scenes recalled as our measure of memory.

### 2.2 Global brain states

To identify global brain states, we first identified neural state boundaries and subsequently clustered the neural state we obtained into two distinct brain states that reoccur across participants and runs.

Neural states boundaries were identified using the Greedy State Boundary Search (GSBS) algorithm. As input to the GSBS algorithm was the ROI × time matrices that we constructed for each subject and run. Although local, within-ROI single subject data has been shown to be noisy for GSBS to identify reliable neural state boundaries, we have found global whole-brain signals to be robust enough for reliable boundary detection. Supplementary analysis showing the reliability of global single subject GSBS can be found at Supplementary Section 4.2. GSBS is a data-driven method that uses an iterative approach to extract neural states from timeseries data (Geerligs et al., 2021). In placing neural states boundaries and determining the optimal number of states, it aims to maximize within-state neural activity correlations and minimize between-state correlations. The maximum number of neural states was set to half the number of time points in a given run. In addition, finetuning was set to 1, which implies that each neural state boundary can be shifted with one TR after each iteration, in case this improves the fit (Geerligs et al., 2021). Furthermore, we used the improved GSBS-states algorithm, introduced in Geerligs et al. (2022). This implementation can place either one or two boundaries per iteration and results in more reliable neural state identification. For each neural state, we computed the average activity pattern, which was subsequently used as input to a clustering algorithm, aimed at identifying shared global brain states across participants.

After identifying global brain state boundaries for each participant with GSBS, we used “Density-based spatial clustering of applications with noise” (DBSCAN) to cluster global neural state patterns to find unique recurring states (Ester et al., 1996). We choose this approach specifically because it determines the optimal number of states based on input parameters. The input to DBSCAN was the pooled state patterns across all subjects and all runs (separately per dataset). As the distance metric to compare average neural state patterns we used cosine distance. DBSCAN has two input parameters; the minimum number of points required to form a dense region (min-samples), and the maximum distance between two samples for one to be considered as in the neighbourhood of the other (eps). To identify the optimal input parameters for our data we performed a grid search between 0 to 1 cosine distance (for eps) and 2 to 2000 minimum number of points (for min-samples) in 20 steps. We scored each of these clusterings with the Silhouette score and Calinski-Harabasz score, and choose the solution with the optimal scores. Full grid search results can be seen at Supplementary Section 4.2. In case the Silhouette and Calinski-Harabasz scores did not agree, we choose the solution with the fewest outlier points. For both datasets, this resulted in two distinct clusters or global brain states. These were mapped onto the cortex by averaging the activity patterns that belong to each global brain state. The global brain states we identified were highly similar to those found in our previous work (Gözükara et al., 2026) and were labelled TPN and DMN. To create a time series of global state labels, we assigned each TR to the cluster label (0 for DMN, 1 for TPN) of its corresponding state. These timeseries were used in further analyses.

### 2.3 GLM and FIR models

In both datasets, we aimed to estimate the neural responses to the occurrence of event (or scene) boundaries. We were interested both in the hippocampal activity around event boundaries and the change in global brain states that occurs around event boundaries. To this end, we used two different approaches.

First we used a conventional general linear model (GLM) analyses, assuming neural responses follow the canonical hemodynamic response function. Next, we investigated the shape of the hemodynamic responses using finite impulse response (FIR) models.

The predictor of interest in the GLM model was one regressor containing all event onsets, which was convolved with the canonical hemodynamic response function. For this GLM analysis, we used the original, not the adjusted event boundary onset.

For the FIR analysis we created a separate predictor for each time-point in the range of +/- 10 TRs relative to the adjusted event boundary onset. This yielded one beta-value per time-point.

In both the FIR and GLM models, we additionally included drift regressors of no interest to model constant, linear and quadratic effects. The same models was used to predict the hippocampal activity and the binary global brain state timeseries. Both the FIR and GLM models were estimated for each participant and subsequently, the Wilcoxon signed rank test was used to test whether the obtained beta values across participants were significantly different from zero.

To investigate the hippocampal response to event boundaries separately for the TPN and DMN global brain states, we performed additional GLM and FIR analyses. First, we labelled event boundaries based on which global brain state was more dominant around a window of -3 +3 TRs. Specifically, when more than 70% of these timepoints were assigned to one state, we assigned the event boundary to that state (5 or more TRs out of 7). Event boundaries which did not have a clearly dominant state around this window were excluded from this analysis. Next, we re-estimated the GLM and FIR models as specified above to predict the hippocampal signal, using TPN and DMN state event boundaries as separate regressors. To investigate whether the difference in hippocampal response in both states could be explained by the differences in mean hippocampal activity in both states, we repeated these analyses including the global state timeseries as a regressor of no interest.

### 2.4 Cortical Mapping and Surface Projections

In all plotted cortical maps, we projected the results of each ROI onto all the voxels they cover. We mapped the volumetric results on an average inflated cortical surface mesh (fsaverage) for visualization purposes. All volumetric nifti files are also available with the code.

## 3 Results

### 3.1 Global Neural State Segmentation and Labelling

To investigate the relationship between event segmentation, memory, and the hippocampal response, we first needed to establish that reliable global brain states could be identified at the individual participant level. We used a three step approach to segment global neural activity into distinct brain states.

First, we extracted the average brain signal from each of 400 Schaefer ROIs across the cortex. Second, we identified neural states boundaries using the Greedy State Boundary Search (GSBS) algorithm. Third, we clustered the resulting neural state activity patterns into two distinct whole brain states using Density-Based Spatial Clustering of Applications with Noise (Ester et al., 1996). More detail on the results of the parameter search for the clustering can be found in Supplementary Section 4.2. The global brain states we identified were highly similar to the ones we identified in our previous work (Gözükara et al., 2026), with one state centred around the DMN (with high activity in the precuneus, posterior cingulate cortex, angular gyrus, medial prefrontal cortex, anterior temporal cortex, and middle temporal gyrus) and another around the TPN (with high activity in frontal eye fields, intraparietal sulcus, visual cortex, anterior insula, temporoparietal junction, and dorsolateral prefrontal cortex) (Figure 1 a, b, d, and e). This was consistent across both the Sherlock and StudyForrest datasets. These results show that the global brain states that we observed in group-average data in our previous work can also be observed on the level of individual participants.

**Figure 1:**
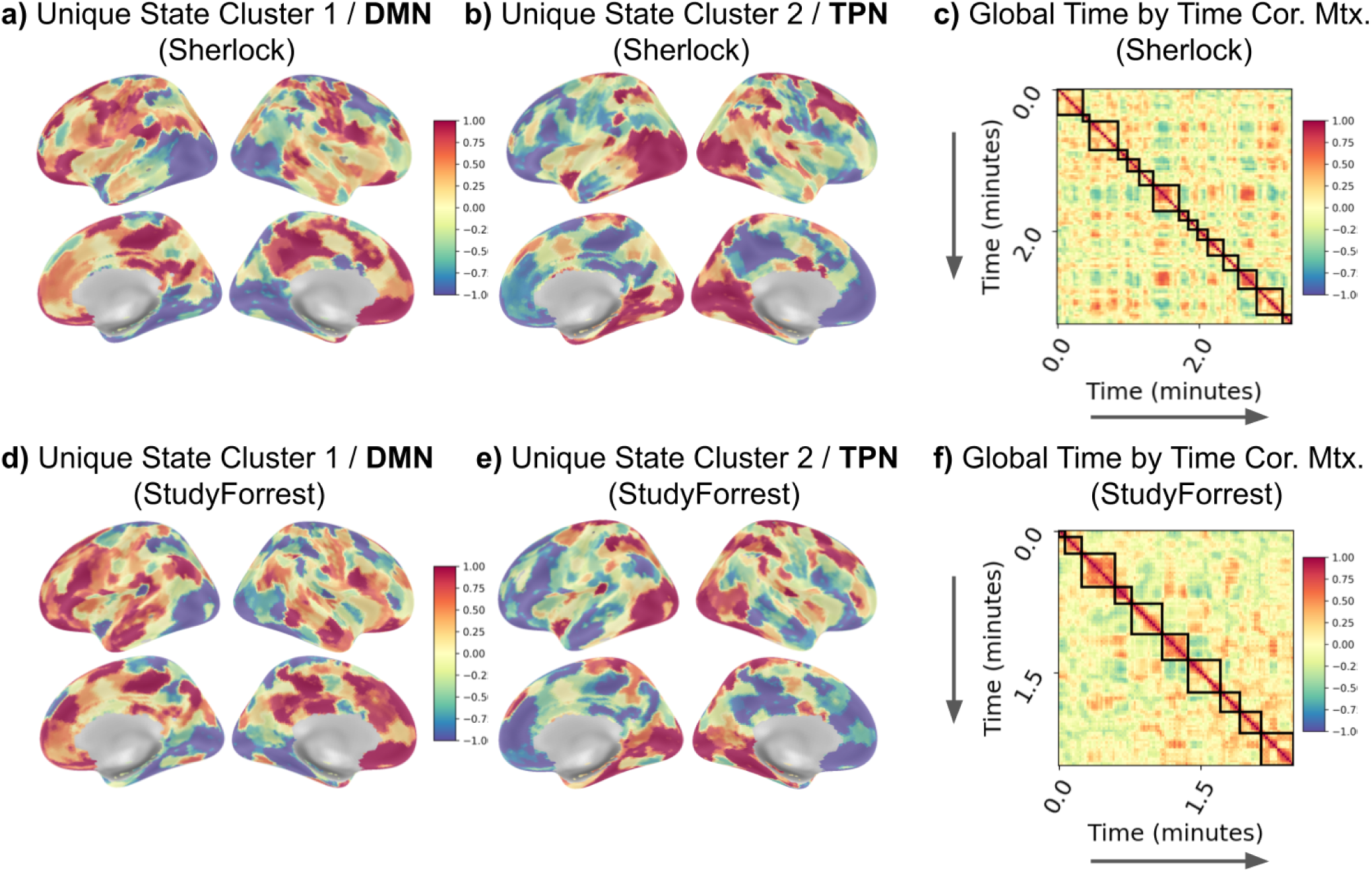
Two unique global brain states, corresponding to Default Mode and Task Positive Networks, are present in movie-watching fMRI data. **Panels (a), (b), (d), (e) - Global Brain Maps:** These panels display the average spatial patterns of the two identified brain state clusters for the **Sherlock dataset (a, b)** and the **StudyForrest dataset (d, e). DMN State (a, d)**: The first cluster map shows a pattern of high activity in areas typically associated with the Default Mode Network (DMN). **TPN State (b, e)**: The second cluster map shows a pattern of high activity in areas typically associated with the Task-Positive Network (TPN). **Panels (c) and (f) - State Boundary Identification**: These panels illustrate the results of Global State Boundary Segmentation (GSBS) on whole-brain fMRI activity. They show the time-by-time Pearson R correlation matrix of an example participant’s Region of Interest (ROI) whole-brain activity for the **Sherlock dataset (c)** and the **StudyForrest dataset (f)**. Black lines indicate the state boundaries automatically detected by the GSBS algorithm.

After clustering, each timepoint was assigned to a cluster label (0 for DMN, 1 for TPN), resulting in a time series of state labels. These timeseries were used in further analyses and GLM and FIR models.

### 3.2 Replicating The Event Boundary Evoked Hippocampal Response

Having established that we can reliably identify global brain states at the individual level, we next sought to replicate the well-documented hippocampal response to event boundaries. This served both as a validation of our pipeline, as well as a foundation for the subsequent state-dependent analyses.

We replicated the previously described event boundary related hippocampal response for both datasets (Figure 2 a and c) using general linear models (GLM) and finite impulse response (FIR) models. For both datasets the GLM analysis revealed a significant increase in mean activity at event boundaries across subjects (StudyForrest: Z=3.40, p*<*0.001; Sherlock: Z=2.86, p=0.005). We then performed a Finite Impulse Response (FIR) analysis to check the timeseries progression around event boundaries (see Figure 2 a and c, right). Both datasets had a statistically significant activity increase around the (HRF-corrected) event boundaries. These results are in line with previous literature (Baldassano et al., 2017; Ben-Yakov & Henson, 2018), and indicate a clear event related activity increase in the hippocampus around event boundaries.

**Figure 2:**
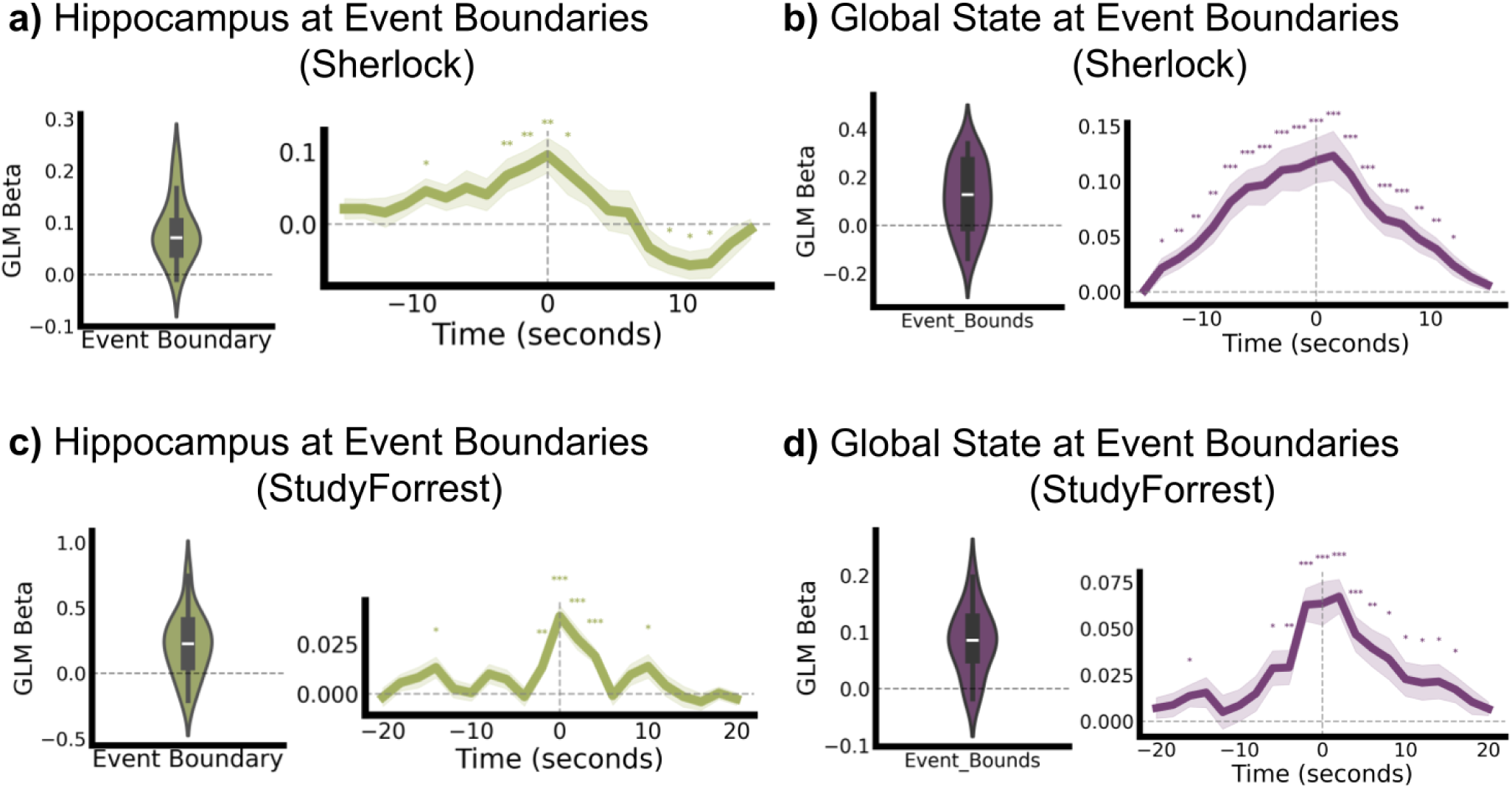
Event boundaries lead to increased hippocampal activity, and a shift to the global TPN brain state. **Panels (a) and (c) - Hippocampal Activity:** These panels show the effect of event boundaries on hippocampal activity, using the **Sherlock dataset (a)** and the **StudyForrest dataset (c)**. **Left (Both Panels):** Violin plots illustrate the distribution of GLM beta coefficients for event boundaries on the hippocampus across all participants. **Right (Both Panels):** Mean FIR beta values across participants are plotted for time points around the event boundaries, which are centred in the graph and adjusted for the expected delay of the Hemodynamic Response Function (HRF; -4.5 seconds). Shaded areas represent the standard error. **Panels (b) and (d) - Global Brain State:** These panels show the effect of event boundaries on the global brain states, using the Sherlock dataset **(b)** and the StudyForrest dataset **(d)**. Because the states are coded such that 0 signified the DMN state and 1 the TPN, a positive beta means higher probability of being at the TPN state, and a negative beta means higher probability of being at the DMN state. **Left (Both Panels):** Violin plots illustrate the distribution of GLM beta coefficients for event boundaries on global brain states across all participants. **Right (Both Panels):** Mean FIR beta values across participants are plotted for time points around the event boundaries, which are centred in the graph and adjusted for the expected delay of the HRF (4.5 seconds). Shaded areas represent the standard error. *Statistical significance:* ^∗^*p<*0.05, ^∗∗^*p<*0.01, ^∗∗∗^*p<*0.001 (vs. baseline)

### 3.3 Replicating The Event Boundary Evoked Global TPN State

In addition to the hippocampal response, we previously observed that event boundaries are associated with a shift toward the TPN state (Gözükara et al., 2026). Here we tested if this association also holds at the individual participant level. To do so, we used GLM and FIR models as the hippocampal activity analysis, but used the binary state label timeseries instead of hippocampal activity (Figure 2b and d). Because the states are coded such that 0 signified the DMN state and 1 the TPN, a positive beta means higher probability of being at the TPN state, and a negative beta means higher probability of being at the DMN state. For both the StudyForrest and Sherlock datasets, the GLM analysis revealed a significant increase in the probability of being in the TPN state after the occurrence of an event boundary (StudyForrest: Z=3.23, p=0.001; Sherlock: Z=2.95, p=0.003). We then performed a Finite Impulse Response (FIR) analysis to check the timeseries progression around event boundaries (see 2). Both datasets had a statistically significant beta estimate increase around the (HRF-corrected) event boundaries. This increased probability started already at 12 seconds before the event boundary for the Sherlock dataset and 18 seconds for the StudyForrest data. And it continued up to 10 seconds after onset for Sherlock and 16 seconds for StudyForrest. These results suggest that participants may be able to anticipate an event boundary coming up and that the increased probability of being in the TPN state occurs for a longer period of time than the increase in hippocampal activity. That being said, it is also possible that the degree of temporal extension could have been exaggerated due to GSBS’s method of state assignment. GSBS partitions the timeseries into discrete states by maximising within-state homogeneity (Geerligs et al., 2021). Timepoints at which the global activity pattern is ambiguous or transitional will therefore be absorbed into whichever neighbouring state they most closely resemble. If the TPN state is particularly pronounced at the moment of an event boundary, weakly TPN-like timepoints preceding it may be drawn into the same state, causing the TPN label to extend further back in time than the underlying neural dynamics would warrant. This would produce an apparently early onset and late offset of the TPN signal in the FIR timecourse that reflects the boundaries of the GSBS state rather than the true timing of the state transition.

### 3.4 Global State Occurrences and Hippocampal Activity

Next, we investigated the activity of the hippocampus in the two different brain states. Figure 3 shows some general descriptives of the temporal characteristics of unique whole brain states during movie-watching, as well as general hippocampal activity during these states. Figure 3a shows that participants spend around 60 percent of their movie watching time at the DMN state, and the remaining 30 at the TPN state. These numbers are consistent across participants, as well as datasets, where the time spent at DMN is statistically significantly higher than at TPN (Wilcoxon signed-rank test, Sherlock: z=3.57, p*<*0.001, StudyForrest: z=3.40, p*<*0.001). Figure 3b shows that mean hippocampal activity is consistently higher at the global TPN state compared to the DMN state for both datasets (Wilcoxon signed-rank test, Sherlock: z=3.62, p*<*0.001, StudyForrest: z=3.40, p*<*0.001). This means that even though the literature consistently shows hippocampal activity to be correlated with ROIs and (sub)networks associated with the DMN (Alves et al., 2019; Andrews-Hanna et al., 2010; Buckner et al., 2008; Ezama et al., 2021; Greicius et al., 2004); the activity of the hippocampus is higher when the brain is in a TPN state. Note that the global states are determined solely by the activity patterns of ROIs that belong to the neocortex, and thus the hippocampal activity did not influence the definition of these states.

**Figure 3:**
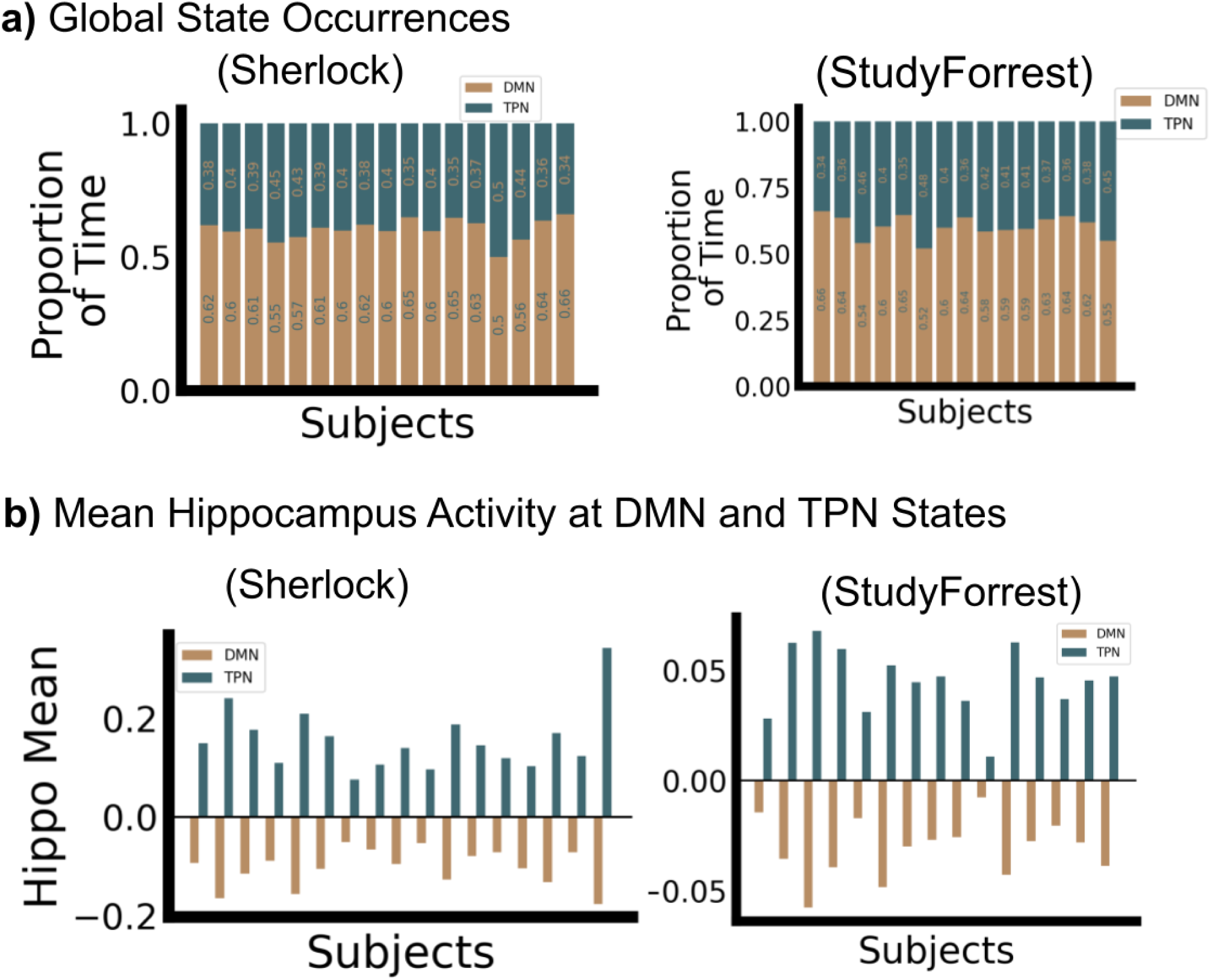
Global brain state occurrences and associated mean hippocampal activity across subjects, using the Sherlock and StudyForrest fMRI datasets. **(a) Global State Occurrences:** Stacked bar charts show the proportion of time each subject spent in the DMN (brown) and TPN (teal) global brain states during the respective stimulus presentation (for Sherlock on the left and for StudyForrest on the right). **(b) Mean Hippocampus Activity at DMN and TPN States:** Bar charts illustrate the mean z scored hippocampus activity during periods when the brain was classified as being in the DMN (brown) or the TPN (teal) state (for Sherlock on the left and for StudyForrest on the right).

### 3.5 Event Boundary Evoked Hippocampal Response at DMN and TPN States

After seeing the consistency of the temporal dynamics of global neural states across participants and datasets, and the consistently higher hippocampal activity in the TPN state, we wanted to see if the hippocampal response to event boundaries is dependent on the global brain states. We labelled event boundaries based on which global brain state was more dominant around a window of -3 +3 timepoints, with a threshold of passing 70%, individually for each participant. Event boundaries which did not have a clearly dominant state around this window were excluded from further analysis (for Sherlock a mean of 10% of all boundaries with a standard deviation of 4%, and for StudyForrest a mean of 17% with a standard deviation of 4%.).

Violin plots in Figure 4 (a) and (b) show that the TPN-labelled boundaries have a distribution of higher GLM beta estimates compared to the DMN-labelled boundaries. For the Sherlock dataset (Figure 4(a)), event boundaries that occurred within a DMN state did not show any significant increase in hippocampal activity (z=0.92, p=0.355), while event boundaries in a TPN state did show this increase (z=3.24, p=0.001). Indeed, the difference in the GLM beta values for the DMN and TPN event boundaries was statistically significant (z=2.34, p=0.019). The same pattern was observed in the StudyForrest dataset. Event boundaries within the DMN state did not show any significant increase in hippocampal activity (z=0.68 p=0.495), while event boundaries within a TPN state did show this increase (z=3.40, p*<*0.001). Again, the difference in the GLM beta values for the DMN and TPN event boundaries was statistically significant (z=3.23, p=0.001). This suggests that the evoked hippocampal response at event boundaries occurs primarily in the TPN state.

**Figure 4:**
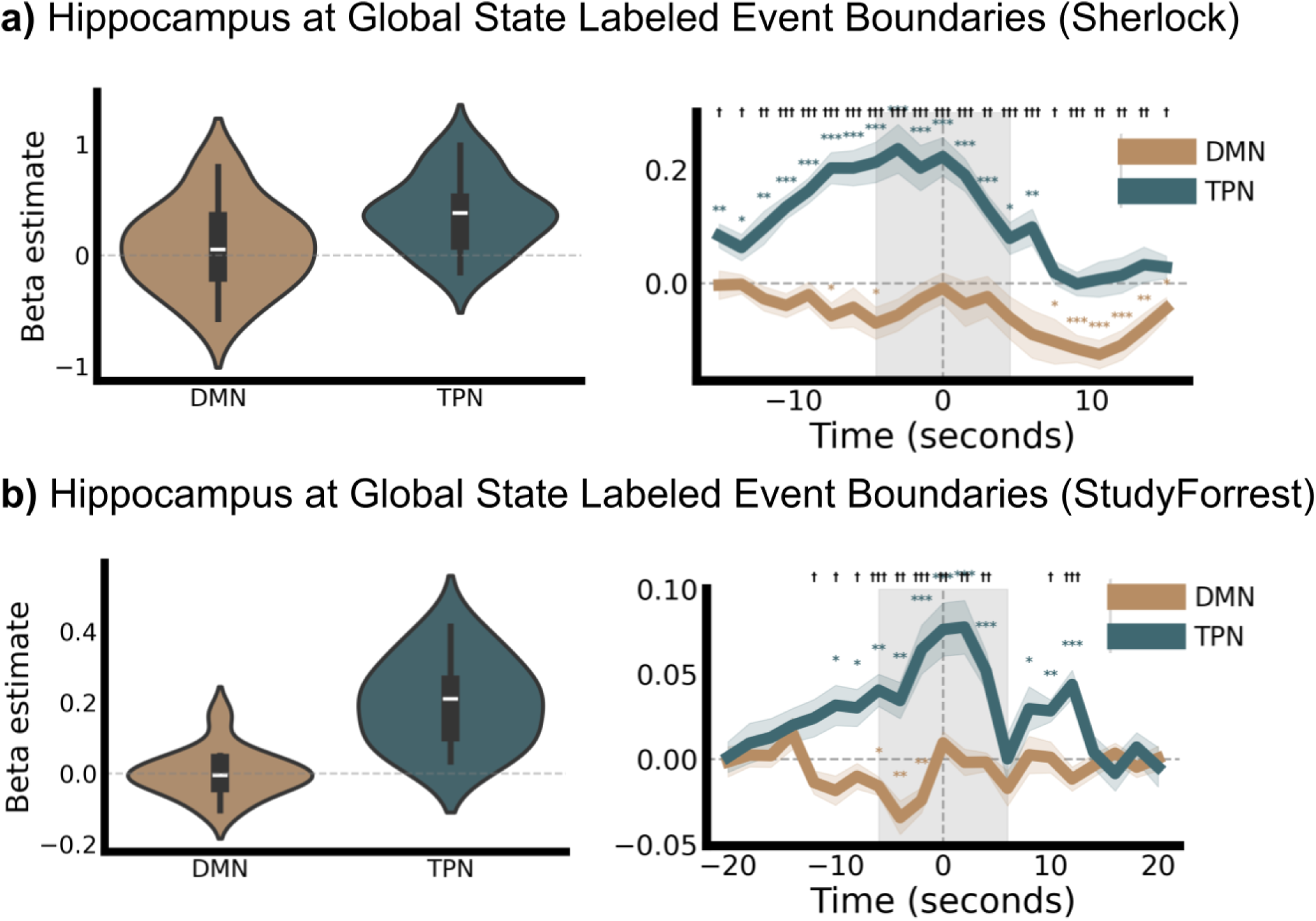
Hippocampal Activity Around Global State-Labelled Event Boundaries for a) Sherlock and b) StudyForrest. **Left panels**: violin plot compares the overall distribution of hippocampal beta estimates (activity levels) during DMN-labelled boundaries (brown) versus TPN-labelled boundaries (teal). The TPN-labelled boundaries show a distribution of hippocampal activity that is shifted toward higher, positive beta estimates compared to the DMN-labelled boundaries. **Right panels**: time course plot shows the mean FIR betas in a time window around the event boundary (time = 0), separately for DMN (brown) and TPN (teal) boundaries. Shaded areas around the lines represent the standard error. The gray shaded area highlights the window used to determine global state label. *Statistical significance:* ^∗^*p<*0.05, ^∗∗^*p<*0.01, ^∗∗∗^*p<*0.001 (vs. baseline); ^†^*p<*0.05, ^††^*p<*0.01, ^†††^*p<*0.001 (between conditions).

To be able to see the timecourse of neural activity in response to event boundaries in different states in more detail, we also ran a FIR beta timeseries on Figure 4(a) and (b). For the Sherlock dataset, Figure 4(a), hippocampal activity is significantly positive during the TPN-labelled boundaries, starting as early as 12 seconds before the event boundary, and falls down to baseline several seconds after the event. In contrast, activity remains near or slightly below zero for the DMN-labelled boundaries, with a statistically insignificant increase at the event boundaries. For the StudyForrest dataset, Figure 4(b), hippocampal activity is significantly positive around the TPN-labelled boundaries, starting around 10 seconds before the (HRF adjusted) event boundary and falls down to baseline only after 12 seconds. In contrast, activity is statistically significantly below zero for the DMN-labelled boundaries pre-boundary, with a return to baseline at the event boundaries.

Taken together, these results indicate that the hippocampal activity increase at event boundaries pre-dominantly occurs when the brain is in the TPN state. However, since hippocampal activity is generally higher during TPN states regardless of event boundaries, this pattern could reflect the overall state-related difference rather than a genuine interaction between brain state and event boundary processing. To disentangle these possibilities, we repeated the analysis while controlling for the overall difference in hippocampal activity between states.

We added the binary global state label timeseries as a covariate to the GLM and FIR analyses, thereby accounting for the overall difference in hippocampal activity between the DMN and TPN states. Once we added this covariate in the GLM, the pattern of results changed in an informative way.

In the Sherlock dataset there was still no significant evoked hippocampal response for event boundaries in the DMN state (z=1.53, p=0.123) and there was a significant response in the TPN state (z=2.24, p=0.024). However, DMN and TPN event boundary GLM beta distributions no longer statistically significantly differed from one another z=0.30, p=0.758). In contrast, even with this covariate, the FIR analysis showed that the hippocampal activity increased around event boundaries at the TPN state, but not in the DMN state (see Figure 5) and that the difference between the two states was still statistically significant. The difference between the GLM and FIR results might be because the hippocampal activity around TPN event boundaries already starts around 12 seconds before the event boundary. Together, these results suggest that in the Sherlock dataset, the hippocampal response is modulated by the global state, even when we use the global state as a covariate. In DMN event boundaries, there is no significant hippocampal response.

**Figure 5:**
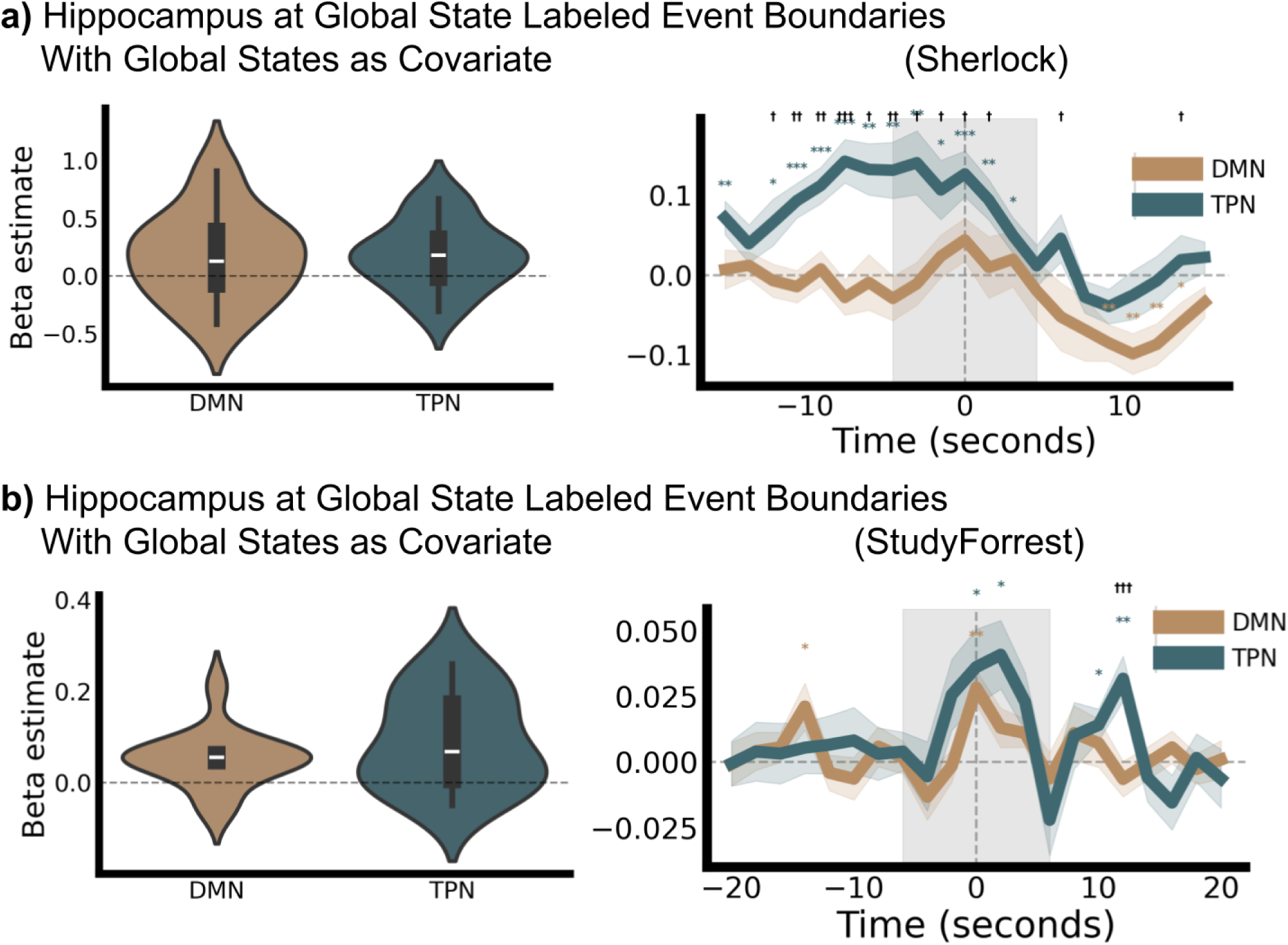
Hippocampal Activity Around Global State-Labelled Event Boundaries. Same as Figure 4 but with global state labels as a covariate in the GLM and FIR models, for a) Sherlock and b) StudyForrest. **Left**: Violin Plot compares the overall distribution of hippocampal beta estimates (activity levels) during DMN-labelled boundaries (brown) versus TPN-labelled boundaries (teal). **Right**: Time course plot shows the mean FIR betas in a time window around the event boundary (time = 0), separately for DMN (brown) and TPN (teal) boundaries. Shaded areas around the lines represent the standard error. The gray shaded area highlights the window used to determine global state label. *Statistical significance:* ^∗^*p <*0.05, ^∗∗^*p<*0.01, ^∗∗∗^*p<*0.001 (vs. baseline); ^†^*p<*0.05, ^††^*p<*0.01, ^†††^*p<*0.001 (between conditions).

For the StudyForrest dataset, Figure 5(a), with the global state labels as a covariate, a different picture emerges. Event boundaries in both DMN and TPN states showed a significant hippocampal evoked response (DMN: z=2.15, p=0.030; TPN: z=2.55, p=0.010). DMN and TPN event boundary GLM beta distributions were no longer statistically significantly differed from one another (z=0.96, p=0.334). Similarly, in the FIR analysis, both DMN and TPN state event boundaries showed a significant increase in hippocampal activity, which was centred narrowly around the HRF-adjusted event boundary. We did not observe a significant difference between TPN and DMN event boundaries in the period around stimulus onset.

These results indicate that when we account for the fact that hippocampal activity is generally higher at the TPN state, we can see that event boundaries, independent of the global brain state, lead to an increase in hippocampal activity for the StudyForrest, but not for the Sherlock dataset. In the Sherlock dataset, we see that TPN event boundaries consistently result in a stronger hippocampal response. While trying to reconcile these results it is worth noting that the StudyForrest dataset is about twice as long in duration than the Sherlock dataset, and has about three times more event boundaries. Therefore we believe the StudyForrest results to be more representative of the general population.

### 3.6 Relation to Memory

Finally, having characterized the temporal dynamics of global brain states and their relationship to hippocampal event boundary responses, we examined how these neural measures relate to subsequent memory performance. This analysis was performed on the Sherlock dataset, for which memory scores were available.

In order to understand how the hippocampal activity and global brain states relate to memory, we checked the correlations between memory scores and the overall time spent in different states, the state occurrence around event boundaries, and the hippocampal response to event boundaries(see Figure 6). The results in Figure 6(a) suggest that there may be a relationship between overall time spent in the DMN state and memory. However, this relationship is not significant (r=0.32, p=0.200) due to one participant who spent much less time in the DMN state than all other participants. If this participant (2.7 standard deviations away from the mean, outlier using the 1.5 IQR method) is removed (see red dot in Figure 6(a)), we do see a significant relationship between time spent in the DMN state and memory (r=0.59, p=0.016). Since there is a direct inverse relationship between the time spent in DMN and the time spent in TPN, this also means that less time spent in TPN is generally associated with better memory. We additionally see in Figure 6(b) that there is a clear positive correlation between being in the TPN state at event boundaries and memory (Pearson R: 0.70, p-value: 0.002), but there is no relationship between overall memory score and the hippocampus activity at event boundaries (Figure 6(c)) (Pearson R: -0.18, p-value: 0.480).

**Figure 6:**
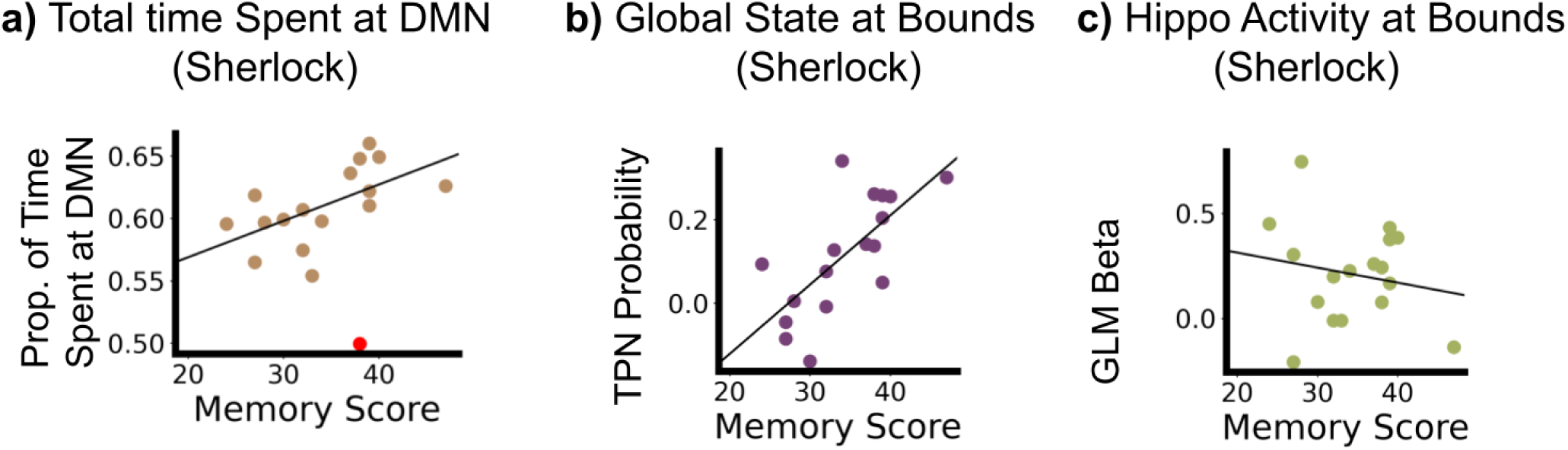
Relationship Between Memory Score and Global State and Hippocampal Activity in the Sherlock Dataset. **a) Memory and Overall DMN Time**: There is a positive correlation between the proportion of time a subject spent in the DMN and their memory score. Red dot denotes outlier subject (2.7 standard deviations away from the mean, outlier using the 1.5 IQR method). Subjects who spent a greater fraction of time in the DMN state tended to have better memory for the movie events. **b) Memory and Global State at Boundaries**: Subjects’ tendency to be in the TPN state at event boundaries plotted against their memory score. The values indicate how much more (or less) likely was that subject’s brain to be in the TPN (task-positive network) state compared to their own personal baseline at the exact moment of an event boundary, showing a positive correlation. Subjects with a stronger tendency to shift into the TPN state at event boundaries showed better subsequent memory. **c) Memory and Hippocampus Activity at Boundaries**: GLM beta coefficient for hippocampus activity at event boundaries against memory scores. This measures the change in hippocampal activity around the moment of an event boundary. No clear relationship is observed between the hippocampal response at event boundaries and subsequent memory performance.

These results suggest that while better memory may be associated with more overall time spent in the DMN state, it is important to switch to the TPN state at the time of an event boundary for better memory. This suggests that the global brain state might be more important (or more reliably measured) than univariate hippocampal activity for memory processes, but further empirical investigation is needed for a clearer judgement.

## 4 Discussion

While it is clear that event boundaries play a critical role in memory formation, the neural underpinnings of this role are only partly know. Previous research has suggested that hippocampal activity at event boundaries reflects encoding of the just-finished event into long-term memory and the online retrieval of relevant past information from previous events (Baldassano et al., 2017; Ben-Yakov & Dudai, 2011; Ben-Yakov & Henson, 2018). Here, we replicate these findings and show that event boundaries also trigger a shift into the TPN state. This shift relates to memory in several ways: (1) the TPN state is generally accompanied by increased hippocampal activity; (2) being in the TPN state around event boundaries predicts better memory; and (3) hippocampal responses to event boundaries may be stronger in the TPN than in the DMN state. Together, these results raise important questions about the role of global attention in event segmentation and memory encoding.

The TPN identified in our analyses is similar to that found in our previous work (Gözükara et al., 2026) and is characterized by heightened engagement of sensory, attention, and executive control networks (Barber et al., 2017; Cheng et al., 2020; Fox et al., 2005; Hammer et al., 2024; Kucyi et al., 2020; Leech et al., 2025; Raichle et al., 2001; Weigard et al., 2024). In our previous work (Gözükara et al., 2026), we found that the TPN state was more prevalent immediately following perceptual changes in naturalistic stimuli (e.g., visual scene changes, auditory transitions) and event boundaries. This is in line with the current results, supporting the role of the TPN in processing salient changes in the environment. The areas that make up the TPN are preferentially engaged during tasks requiring focused attention, working memory, and cognitive control (Barber et al., 2017; Cheng et al., 2020; Fox et al., 2005; Hammer et al., 2024; Kucyi et al., 2020; Leech et al., 2025; Raichle et al., 2001; Weigard et al., 2024). Increased TPN activity at event boundaries may therefore reflect increased externally-directed cognition and increase attention toward new perceptual input. Indeed, event segmentation has been linked to a transient increase in attention (Radvansky & Zacks, 2017). Describing the TPN as a state of externally directed processing or externally directed attention is also in line with previous research on the frontal eye fields (FEF) and intraparietal sulcus (IPS). These areas are a core part of the TPN and form a bilateral frontoparietal circuit that governs the goal-directed deployment of visuospatial attention(Corbetta & Shulman, 2002; Fox et al., 2005), suggesting that event boundaries recruit attentional mechanisms that facilitate the processing of narrative transitions.

Such attentional changes at event boundaries are likely to play an important role in memory formation, as suggested by our observation that higher TPN state probability is related to better memory and a stronger hippocampal response. This link between attention and memory encoding is well established, where attention during encoding is a critical determinant of what is subsequently remembered. Classic divided-attention studies have demonstrated that splitting attentional resources during encoding produces large and consistent reductions in later memory performance, whereas division of attention at retrieval has comparatively little effect (Chun & Turk-Browne, 2007; Naveh-Benjamin et al., 1998) At the neural level, activation in dorsal parietal regions, particularly the IPS and superior parietal lobule, during encoding predicts later memory success, linking the engagement of the dorsal attention system directly to the formation of durable memory traces (Uncapher & Wagner, 2009). More recently, Aly and Turk-Browne (2016) demonstrated that behavioural goals modulate the attentional state of the hippocampus itself, stabilising task-relevant representations and thereby prioritising what aspects of an experience are encoded. These findings indicate that the attentional mechanisms instantiated by the TPN are important for determining what information enters long-term memory.

When we investigated the association with memory recall, we found that participants who spent more overall time in the DMN state and importantly who also showed a more pronounced shift toward the TPN state specifically at event boundaries had better memory. This highlights the importance of timely attentional engagement at moments of narrative change. Spending more time overall in the DMN state may reflect a processing mode conducive to sustained narrative comprehension, schema maintenance, and the integration of incoming information with existing knowledge structures; processes that the DMN has been proposed to support during naturalistic cognition (Buckner et al., 2008; Menon, 2023). At the same time, the selective shift toward the TPN at event boundaries suggests that timely attentional engagement at moments of narrative change is critical for successful encoding. In other words, an optimal pattern for memory may involve predominantly DMN-driven processing for ongoing narrative understanding, punctuated by transient TPN engagement when the situation model needs to be updated. This is consistent with the broader principle that the dynamic interplay between these anti-correlated networks supports flexible cognition (Fox et al., 2005; Gözükara et al., 2026).

When we look more specifically at the hippocampal response, we observed two things. First, the hippocampus was generally more active in the TPN than DMN states. This is surprising, as the hippocampus is typically considered part of the extended DMN based on resting-state functional connectivity (Ranganath & Ritchey, 2012; Rugg & Vilberg, 2013). Second, we found that the hippocampal response to event boundaries was higher in the TPN state than the DMN state. Once we controlled for the overall differences in hippocampal activity between states, this was still the case for one of the two datasets. This has implications for how the existing event boundary literature on hippocampal responses should be interpreted (Baldassano et al., 2017; Ben-Yakov & Henson, 2018). Given that the hippocampal response at event boundaries is substantially larger when the brain is in a TPN state, the signals reported in that literature likely reflect, at least in part, fluctuations in hippocampal activity that are associated with global state transitions. Therefore, it is not clear to what extent the previously observed hippocampal responses purely reflect an encoding response tied to the occurrence of an event boundary. Including global brain state as a covariate of interest in future studies on hippocampal responses could disentangle effects of state-driven activity changes from the effects of event-specific encoding processes.

Interestingly, the magnitude of the hippocampal response at event boundaries was not correlated with memory performance. Although this could well be due to the small sample size in this data, it might also suggest that global brain state dynamics may be more informative of memory outcomes than univariate hippocampal activity alone. Another possibility is that increased hippocampal activity does not necessarily relate to better memory. Previous work in other cognitive domains has shown that better performance may be accompanied by greater neural efficiency and therefore reduced brain activity (Bernardi et al., 2013; Haier et al., 1988; Krings et al., 2000; Neubauer & Fink, 2009; Rypma et al., 2002). In this context, the elevated hippocampal activity in the TPN state could reflect effortful detection of a narrative change or an attentional orienting response, rather than encoding success per se, and the memory advantage associated with TPN boundaries may be driven by mechanisms that are orthogonal to hippocampal activity levels. Disentangling these possibilities would benefit from approaches that go beyond univariate activity measures. For example, multivariate pattern analyses of hippocampal reinstatement during naturalistic encoding have shown that the degree to which hippocampal activity patterns are reactivated at recall predicts memory for narrative events independently of univariate activation levels (Baldassano et al., 2017; Cohn-Sheehy et al., 2021; Hahamy et al., 2023; Silva et al., 2019). Applying these multivariate analyses to the state-dependent responses we observe would allow a cleaner dissociation between hippocampal activity that reflects effortful event detection and activity that reflects the binding of event representations into long-term memory.

### 4.1 Limitations and Future Directions

Several limitations of the present study should be noted. First, the relationship between global brain states and memory was assessed only in the Sherlock dataset, and the sample size was modest (*N* = 17). The suggestive finding that overall DMN time predicts memory depended on the exclusion of a single outlier. Future studies with larger samples are needed to more definitively establish how the temporal dynamics of global brain states relate to individual differences in naturalistic memory.

Second, while our findings suggest that attention plays a role in the TPN-associated hippocampal response at event boundaries, we did not directly measure attention. Integrating our approach with a divided attention task (as in Naci et al. (2014)) would allow for a more direct test of whether the TPN state at event boundaries reflects attentional engagement.

Finally, future work should examine the role of hippocampal subfield contributions and the directionality of hippocampal-cortical interactions at event boundaries. The hippocampus is not a monolithic structure, and different subfields may contribute differentially to encoding versus retrieval processes at event boundaries (Adnan et al., 2016; Bakker et al., 2008; Barnett et al., 2019; Blessing et al., 2016; Chase et al., 2015; Dalton et al., 2019; Ezama et al., 2021; Fanselow & Dong, 2010). Similarly, effective connectivity analyses could reveal whether the hippocampus drives or is driven by global brain state transitions at event boundaries, providing mechanistic insight into the relationship between large-scale neural dynamics and hippocampal memory processes.

### 4.2 Conclusion

In summary, our study demonstrates that hippocampal responses to event boundaries during naturalistic movie-watching are modulated by the concurrent global brain state. While hippocampal activity is generally elevated during TPN states, event boundaries elicit an independent hippocampal response that is present regardless of the global state once baseline differences are accounted for. The association between the TPN state at event boundaries and subsequent memory, suggests that the brain’s large-scale network dynamics play a critical role in determining when and how continuous experience is segmented and encoded into memory.

## Data and Code Availability

All code required to reproduce the analyses reported in this paper is publicly available at https://github.com/drgzkr/global-states-event-memory.

## Author Contributions

**Dora Gözükara**: Conceptualization, Methodology, Software, Formal analysis, Visualization, Writing - Original Draft. **Nasir Ahmad**: Methodology, Software, Writing - Review & Editing, Supervision. **Djamari Oetringer**: Methodology, Data Pre-processing, Software, Writing - Review & Editing. **Linda Geerligs**: Conceptualization, Methodology, Resources, Writing - Review & Editing, Supervision.

## Acknowledgements

We would like to thank Sahel Azizpour for fruitful discussions throughout the development of this paper.

## Supplementary Material

### Reliability of Whole-Brain GSBS Segmentation

To verify that the global neural state boundaries identified at the individual participant level were reliable and not driven by noise, we assessed the inter-subject consistency of the GSBS segmentation results using a leave-one-out (LOO) correlation approach. For each participant and each run, we extracted the boundary delta vector produced by GSBS (a timeseries vector of 0s and 1s where 1s indicate a neural state boundary) We then computed the mean delta vector across all other participants and calculated the Pearson correlation between participant’s delta vector and this LOO mean. This procedure was repeated for all participants and all runs; per-participant reliability estimates were subsequently averaged across runs.

**Figure 7:**
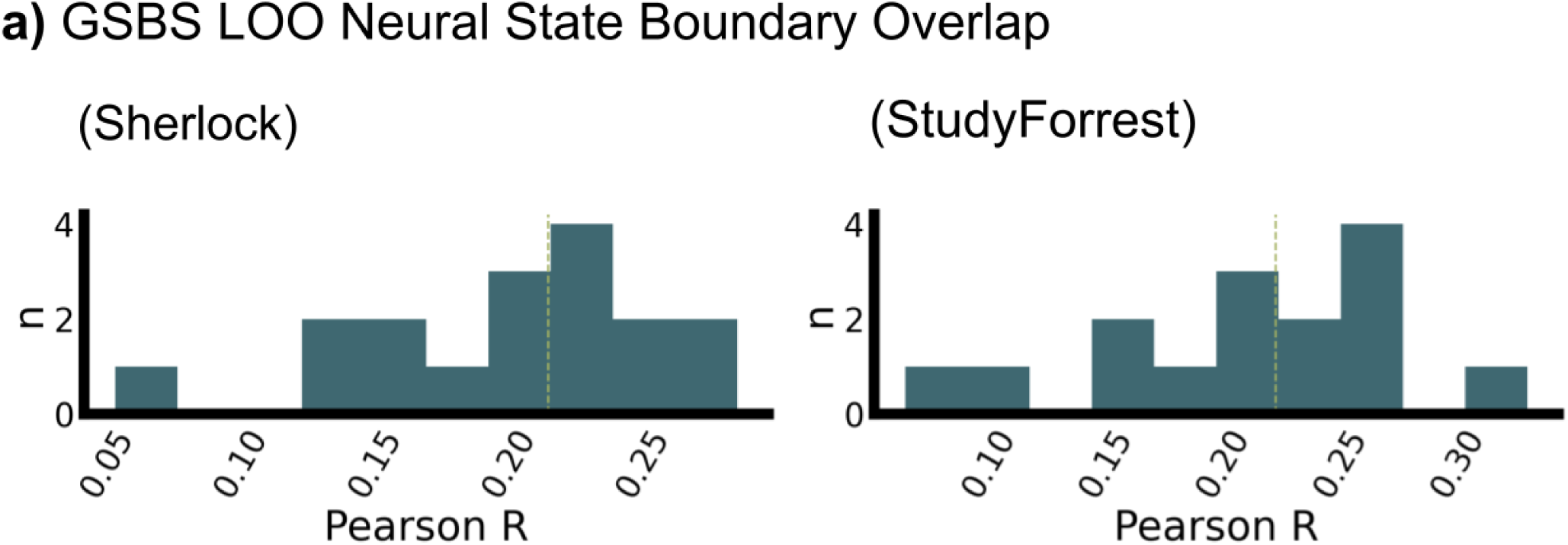
Reliability of Whole-Brain GSBS Neural State Boundaries Across Participants. Histogram of leave-one-out (LOO) inter-subject correlations of GSBS boundary delta vectors for the **(a) Sherlock** and **(b) StudyForrest** datasets. For each participant, the boundary delta vector (a vector of 0s and 1s where 1s indicate a boundary) was correlated with the mean delta vector computed from all other participants. Values were averaged across runs per participant. The dashed vertical line indicates the median correlation across participants. A Wilcoxon signed-rank test confirmed that LOO correlations were significantly greater than zero in both datasets (StudyForrest: *Z* = 3.40, *p <* 0.001; Sherlock: *Z* = 3.62, *p <* 0.001), indicating that individual-level GSBS segmentations reliably recover a common boundary structure shared across participants.

The resulting distribution of LOO correlations was tested against zero using the Wilcoxon signed-rank test. In both datasets, LOO correlations were significantly positive (Sherlock: ; StudyForrest: ), indicating that the boundary structure identified in any individual participant was systematically similar to the consensus structure observed across the rest of the group. This demonstrates that the whole-brain GSBS segmentation at the individual level captures a reliable, shared neural signal rather than subject-specific noise, and justifies the use of individual-level state labels in subsequent GLM and FIR analyses.

### DBSCAN Parameter Selection for Global Brain State Clustering

To identify recurring whole-brain activity patterns across participants, we applied Density-Based Spatial Clustering of Applications with Noise (Ester et al., 1996) to the pooled set of GSBS state activity patterns from all participants and all runs within each dataset. DBSCAN was chosen over fixed-k methods such as k-means because it determines the number of clusters directly from the data rather than requiring it to be specified in advance. Cosine distance was used as the dissimilarity metric when comparing state patterns, as it captures differences in the spatial distribution of activity across ROIs independently of overall signal amplitude.

DBSCAN has two free hyperparameters: *eps*, the maximum cosine distance between two points for one to be considered in the neighbourhood of the other, and *min samples*, the minimum number of points required to form a dense region (and thus a cluster). To identify the parameter combination best suited to our data, we performed a grid search over 20 linearly spaced values of *eps* (range: 0.01–1.00) and 20 linearly spaced values of *min samples* (range: 2–2000), yielding 400 candidate solutions per dataset. For each solution, we computed two complementary internal cluster validity indices: the Silhouette score, which measures how well each point matches its assigned cluster relative to neighbouring clusters, and the CalinskiHarabasz score, which assesses the ratio of between-cluster to within-cluster dispersion. Parameter combinations that resulted in fewer than two clusters were excluded from scoring.

The optimal parameter combination was selected as the solution that maximised both indices. In cases where the Silhouette and Calinski-Harabasz scores favoured different solutions, preference was given to the solution with fewer outlier points (i.e., points assigned the noise label −1 by DBSCAN).

For the Sherlock dataset, the Silhouette score was maximised at *eps* = 0.0621, *min samples* = 2, yielding 2 clusters and 1975 outlier points. The Calinski-Harabasz score was maximised at *eps* = 0.8437, *min samples* = 738, also yielding 2 clusters but with no outlier points; this solution was selected as the final clustering for the Sherlock dataset. For the StudyForrest dataset, both the Silhouette and Calinski-Harabasz scores converged on the same solution (*eps* = 0.7916, *min samples* = 1263), yielding 2 clusters and no outlier points.

In both datasets, the two resulting clusters closely replicated the global brain states reported in our previous work (Gözükara et al., 2026), with one cluster corresponding to a Default Mode Network (DMN)-dominant activity pattern and the other to a Task-Positive Network (TPN)-dominant pattern (Figure 1). The convergence on two clusters across both datasets, two independent scoring criteria, and a wide range of tested hyperparameters provides strong support for the two-state solution adopted throughout the main analyses.

---

**Figure 8:**
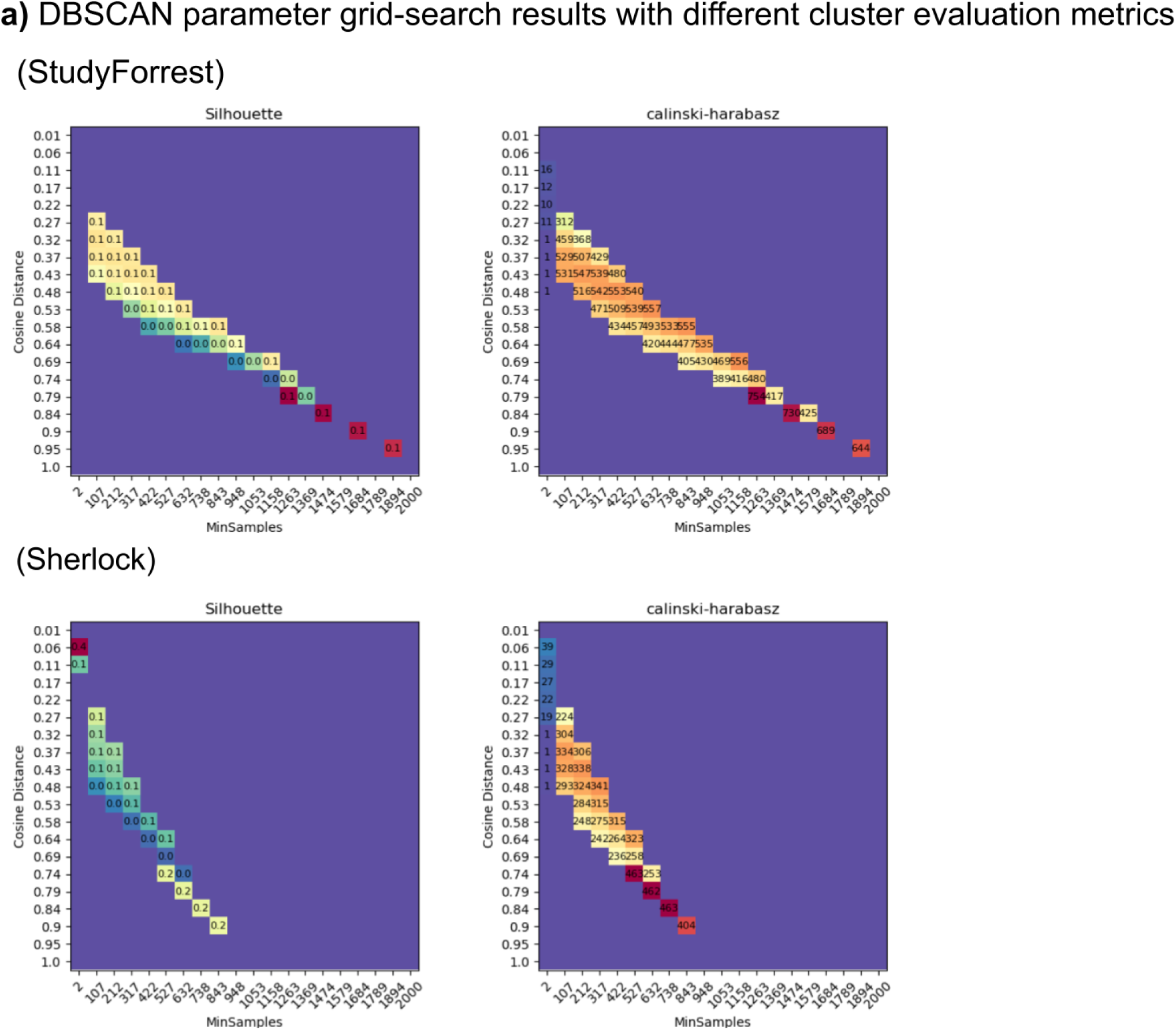
DBSCAN Parameter Grid Search for Global Brain State Clustering. Results of the parameter grid search used to identify the optimal DBSCAN hyperparameters for clustering whole-brain GSBS state patterns into recurring global brain states, shown for the **above, StudyForrest** and **below, Sherlock** datasets. Each panel displays two 20 × 20 heatmaps, with the cosine distance threshold (*eps*; range: 0.01–1.00, in 20 equal steps) on the y-axis and the minimum neighbourhood size (*min samples*; range: 2–2000, in 20 equal steps) on the x-axis. **Left heatmap (Silhouette score):** Silhouette scores for each parameter combination; cells with no valid clustering (fewer than two clusters) are left blank. **Right heatmap (Calinski-Harabasz score):** Calinski-Harabasz scores for each parameter combination; cells with no valid clustering are left blank. Annotated values within each cell indicate the numeric score. The parameter combination maximising each score was selected as the candidate optimal solution; when the two criteria disagreed, the solution with fewer outlier points was preferred. In both datasets, the selected solution yielded two clusters, corresponding to the DMN and TPN global brain states reported in the main text.

## References

Adnan, A., Barnett, A., Moayedi, M., McCormick, C., Cohn, M., & McAndrews, M. P. (2016). Distinct hippocampal functional networks revealed by tractography-based parcellation. Brain Structure and Function, 221 (6), 2999–3012. 10.1007/s00429-015-1084-x

Alves, P. N., Foulon, C., Karolis, V., Bzdok, D., Margulies, D. S., Volle, E., & Thiebaut de Schotten, M. (2019). An improved neuroanatomical model of the default-mode network reconciles previous neuroimaging and neuropathological findings. Communications Biology, 2 (1), 370. 10.1038/s42003-019-0611-3

Aly, M., & Turk-Browne, N. B. (2016). Attention Stabilizes Representations in the Human Hippocampus. Cerebral Cortex, 26 (2), 783–796. 10.1093/cercor/bhv041

Andrews-Hanna, J. R., Reidler, J. S., Sepulcre, J., Poulin, R., & Buckner, R. L. (2010). Functional-Anatomic Fractionation of the Brain’s Default Network. Neuron, 65 (4), 550–562. 10.1016/j.neuron.2010.02.005

Bakker, A., Kirwan, C. B., Miller, M., & Stark, C. E. L. (2008). Pattern Separation in the Human Hippocampal CA3 and Dentate Gyrus. Science, 319 (5870), 1640–1642. 10.1126/science.1152882

Baldassano, C., Chen, J., Zadbood, A., Pillow, J. W., Hasson, U., & Norman, K. A. (2017). Discovering Event Structure in Continuous Narrative Perception and Memory [Publisher: Elsevier]. Neuron, 95 (3), 709–721.e5. 10.1016/j.neuron.2017.06.041

Barber, A. D., Caffo, B. S., Pekar, J. J., & Mostofsky, S. H. (2017). Decoupling of reaction time-related default mode network activity with cognitive demand. Brain Imaging and Behavior, 11 (3), 666–676. 10.1007/s11682-016-9543-4

Barnett, A. J., Man, V., & McAndrews, M. P. (2019). Parcellation of the Hippocampus Using Resting Functional Connectivity in Temporal Lobe Epilepsy. Frontiers in Neurology, 10. 10.3389/fneur.2019.00920

Ben-Yakov, A., & Dudai, Y. (2011). Constructing Realistic Engrams: Poststimulus Activity of Hippocampus and Dorsal Striatum Predicts Subsequent Episodic Memory. Journal of Neuroscience, 31 (24), 9032–9042. 10.1523/JNEUROSCI.0702-11.2011

Ben-Yakov, A., Eshel, N., & Dudai, Y. (2013). Hippocampal immediate poststimulus activity in the encoding of consecutive naturalistic episodes. Journal of Experimental Psychology: General, 142 (4), 1255–1263. 10.1037/a0033558

Ben-Yakov, A., & Henson, R. N. (2018). The Hippocampal Film Editor: Sensitivity and Specificity to Event Boundaries in Continuous Experience [Publisher: Society for Neuroscience Section: Research Articles]. Journal of Neuroscience, 38 (47), 10057–10068. 10.1523/JNEUROSCI.0524-18.2018

Bernardi, G., Ricciardi, E., Sani, L., Gaglianese, A., Papasogli, A., Ceccarelli, R., Franzoni, F., Galetta, F., Santoro, G., Goebel, R., & Pietrini, P. (2013). How Skill Expertise Shapes the Brain Functional Architecture: An fMRI Study of Visuo-Spatial and Motor Processing in Professional Racing-Car and Näıve Drivers. PLOS ONE, 8 (10), e77764. 10.1371/journal.pone.0077764

Blessing, E. M., Beissner, F., Schumann, A., Brünner, F., & Bär, K.-J. (2016). A data-driven approach to mapping cortical and subcortical intrinsic functional connectivity along the longitudinal hippocampal axis [_eprint: https://onlinelibrary.wiley.com/doi/pdf/10.1002/hbm.23042]. Human Brain Mapping, 37 (2), 462–476. 10.1002/hbm.23042

Buckner, R. L., Andrews-Hanna, J. R., & Schacter, D. L. (2008). The brain’s default network: Anatomy, function, and relevance to disease. Annals of the New York Academy of Sciences, 1124, 1–38. 10.1196/annals.1440.011

Chase, H. W., Clos, M., Dibble, S., Fox, P., Grace, A. A., Phillips, M. L., & Eickhoff, S. B. (2015). Evidence for an anterior–posterior differentiation in the human hippocampal formation revealed by meta-analytic parcellation of fMRI coordinate maps: Focus on the subiculum. NeuroImage, 113, 44–60. 10.1016/j.neuroimage.2015.02.069

Chen, J., Leong, Y. C., Honey, C. J., Yong, C. H., Norman, K. A., & Hasson, U. (2017). Shared memories reveal shared structure in neural activity across individuals. Nature Neuroscience, 20 (1), 115–125. 10.1038/nn.4450

Cheng, X., Yuan, Y., Wang, Y., & Wang, R. (2020). Neural antagonistic mechanism between default-mode and task-positive networks. Neurocomputing, 417, 74–85. 10.1016/j.neucom.2020.07.079

Chun, M. M., & Turk-Browne, N. B. (2007). Interactions between attention and memory. Current Opinion in Neurobiology, 17 (2), 177–184. 10.1016/j.conb.2007.03.005

Cohn-Sheehy, B. I., Delarazan, A. I., Reagh, Z. M., Crivelli-Decker, J. E., Kim, K., Barnett, A. J., Zacks, J. M., & Ranganath, C. (2021). The hippocampus constructs narrative memories across distant events. Current Biology, 31 (22), 4935–4945.e7. 10.1016/j.cub.2021.09.013

Corbetta, M., & Shulman, G. L. (2002). Control of goal-directed and stimulus-driven attention in the brain. Nature Reviews. Neuroscience, 3 (3), 201–215. 10.1038/nrn755

Dalton, M. A., McCormick, C., & Maguire, E. A. (2019). Differences in functional connectivity along the anterior-posterior axis of human hippocampal subfields. NeuroImage, 192, 38–51. 10.1016/j.neuroimage.2019.02.066

Ester, M., Kriegel, H.-P., Sander, J., & Xu, X. (1996). A density-based algorithm for discovering clusters in large spatial databases with noise. Proceedings of the Second International Conference on Knowledge Discovery and Data Mining, 226–231.

Ezama, L., Hernández-Cabrera, J. A., Seoane, S., Pereda, E., & Janssen, N. (2021). Functional connectivity of the hippocampus and its subfields in resting-state networks [_eprint: https://onlinelibrary.wiley.com/doi/ European Journal of Neuroscience, 53 (10), 3378–3393. 10.1111/ejn.15213

Fanselow, M. S., & Dong, H.-W. (2010). Are the Dorsal and Ventral Hippocampus Functionally Distinct Structures? Neuron, 65 (1), 7–19. 10.1016/j.neuron.2009.11.031

Fox, M. D., Snyder, A. Z., Vincent, J. L., Corbetta, M., Van Essen, D. C., & Raichle, M. E. (2005). The human brain is intrinsically organized into dynamic, anticorrelated functional networks. Proceedings of the National Academy of Sciences, 102 (27), 9673–9678. 10.1073/pnas.0504136102

Geerligs, L., Gözükara, D., Oetringer, D., Campbell, K. L., Gerven, M. v., & Güçlü, U. (2022, September). A partially nested cortical hierarchy of neural states underlies event segmentation in the human brain [Publisher: eLife Sciences Publications Limited]. 10.7554/eLife.77430

Geerligs, L., van Gerven, M., & Güçlü, U. (2021). Detecting neural state transitions underlying event segmentation. NeuroImage, 236, 118085. 10.1016/j.neuroimage.2021.118085

Gözükara, D., Oetringer, D., Ahmad, N., & Geerligs, L. (2026). Multi-Scale Anti-Correlated Neural States Dominate Naturalistic Whole-Brain Activity. eLife, 15. 10.7554/eLife.109116.1

Greicius, M. D., Srivastava, G., Reiss, A. L., & Menon, V. (2004). Default-mode network activity distinguishes Alzheimer’s disease from healthy aging: Evidence from functional MRI. Proceedings of the National Academy of Sciences, 101 (13), 4637–4642. 10.1073/pnas.0308627101

Hahamy, A., Dubossarsky, H., & Behrens, T. E. J. (2023). The human brain reactivates context-specific past information at event boundaries of naturalistic experiences. Nature Neuroscience, 26 (6), 1080–1089. 10.1038/s41593-023-01331-6

Haier, R. J., Siegel, B. V., Nuechterlein, K. H., Hazlett, E., Wu, J. C., Paek, J., Browning, H. L., & Buchsbaum, M. S. (1988). Cortical glucose metabolic rate correlates of abstract reasoning and attention studied with positron emission tomography. Intelligence, 12 (2), 199–217. 10.1016/0160-2896(88)90016-5

Hammer, J., Kajsova, M., Kalina, A., Krysl, D., Fabera, P., Kudr, M., Jezdik, P., Janca, R., Krsek, P., & Marusic, P. (2024). Antagonistic behavior of brain networks mediated by low-frequency oscillations: Electrophysiological dynamics during internal–external attention switching. Communications Biology, 7 (1), 1105. 10.1038/s42003-024-06732-2

Hanke, M., Adelhöfer, N., Kottke, D., Iacovella, V., Sengupta, A., Kaule, F. R., Nigbur, R., Waite, A. Q., Baumgartner, F., & Stadler, J. (2016). A studyforrest extension, simultaneous fMRI and eye gaze recordings during prolonged natural stimulation [Publisher: Nature Publishing Group]. Scientific Data, 3 (1), 160092. 10.1038/sdata.2016.92

Hanke, M., Baumgartner, F. J., Ibe, P., Kaule, F. R., Pollmann, S., Speck, O., Zinke, W., & Stadler, J. (2014). A high-resolution 7-Tesla fMRI dataset from complex natural stimulation with an audio movie. Scientific Data, 1 (1), 140003. 10.1038/sdata.2014.3

Henderson, S., Oetringer, D., Geerligs, L., & Campbell, K. L. (2025, January). Neural state changes during movie watching relate to episodic memory in younger and older adults. 10.31234/osf.io/pwkht

Kaefer, K., Stella, F., McNaughton, B. L., & Battaglia, F. P. (2022). Replay, the default mode network and the cascaded memory systems model. Nature Reviews. Neuroscience, 23 (10), 628–640. 10.1038/s41583-022-00620-6

Krings, T., Töpper, R., Foltys, H., Erberich, S., Sparing, R., Willmes, K., & Thron, A. (2000). Cortical activation patterns during complex motor tasks in piano players and control subjects. A functional magnetic resonance imaging study. Neuroscience Letters, 278 (3), 189–193. 10.1016/S0304-3940(99)00930-1

Kucyi, A., Daitch, A., Raccah, O., Zhao, B., Zhang, C., Esterman, M., Zeineh, M., Halpern, C. H., Zhang, K., Zhang, J., & Parvizi, J. (2020). Electrophysiological dynamics of antagonistic brain networks reflect attentional fluctuations. Nature Communications, 11 (1), 325. 10.1038/s41467-019-14166-2

Kurby, C. A., & Zacks, J. M. (2008). Segmentation in the perception and memory of events. Trends in cognitive sciences, 12 (2), 72–79. 10.1016/j.tics.2007.11.004

Leech, R., Braga, R. M., Haydock, D., Vowles, N., Jefferies, E., Bernhardt, B., Turkheimer, F., Alberti, F., Margulies, D., Sherwood, O., Jones, E. J., Smallwood, J., & Váša, F. (2025). The spatial layout of antagonistic brain regions is explicable based on geometric principles. Communications Biology, 8 (1), 889. 10.1038/s42003-025-08295-2

Liu, X., Zhen, Z., Yang, A., Bai, H., & Liu, J. (2019). A manually denoised audio-visual movie watching fMRI dataset for the studyforrest project. Scientific Data, 6 (1), 295. 10.1038/ s41597-019-0303-3

Maldjian, J. A., Laurienti, P. J., Kraft, R. A., & Burdette, J. H. (2003). An automated method for neuroanatomic and cytoarchitectonic atlas-based interrogation of fMRI data sets. NeuroImage, 19 (3), 1233–1239. 10.1016/S1053-8119(03)00169-1

Menon, V. (2023). 20 years of the default mode network: A review and synthesis. Neuron, 111 (16), 2469–2487. 10.1016/j.neuron.2023.04.023

Naci, L., Cusack, R., Anello, M., & Owen, A. M. (2014). A common neural code for similar conscious experiences in different individuals. Proceedings of the National Academy of Sciences, 111 (39), 14277–14282. 10.1073/pnas.1407007111

Naveh-Benjamin, M., Craik, F. I. M., Guez, J., & Dori, H. (1998). Effects of divided attention on encoding and retrieval processes in human memory: Further support for an asymmetry. Journal of Experimental Psychology: Learning, Memory, and Cognition, 24 (5), 1091–1104. 10.1037/0278-7393.24.5.1091

Neubauer, A. C., & Fink, A. (2009). Intelligence and neural efficiency. Neuroscience & Biobehavioral Reviews, 33 (7), 1004–1023. 10.1016/j.neubiorev.2009.04.001

Oetringer, D., Gözükara, D., Güçlü, U., & Geerligs, L. (2024, November). The Neural Basis of Event Segmentation: Stable Features in the Environment are Reflected by Neural States [Pages: 2024.01.26.577369 Section: New Results]. 10.1101/2024.01.26.577369

Oetringer, D., Gözükara, D., Güçlü, U., & Geerligs, L. (2025). The neural basis of event segmentation: Stable features in the environment are reflected by neural states. Imaging Neuroscience, 3. 10.1162/imag_a_00432

Radvansky, G. A., & Zacks, J. M. (2017). Event boundaries in memory and cognition. Current Opinion in Behavioral Sciences, 17, 133–140. 10.1016/j.cobeha.2017.08.006

Raichle, M. E., MacLeod, A. M., Snyder, A. Z., Powers, W. J., Gusnard, D. A., & Shulman, G. L. (2001). A default mode of brain function. Proceedings of the National Academy of Sciences, 98 (2), 676–682. 10.1073/pnas.98.2.676

Ranganath, C., & Ritchey, M. (2012). Two cortical systems for memory-guided behaviour [Publisher: Nature Publishing Group]. Nature Reviews Neuroscience, 13 (10), 713–726. 10.1038/nrn3338

Reagh, Z. M., Delarazan, A. I., Garber, A., & Ranganath, C. (2020). Aging alters neural activity at event boundaries in the hippocampus and Posterior Medial network. Nature Communications, 11 (1), 3980. 10.1038/s41467-020-17713-4

Rugg, M. D., & Vilberg, K. L. (2013). Brain networks underlying episodic memory retrieval. Current Opinion in Neurobiology, 23 (2), 255–260. 10.1016/j.conb.2012.11.005

Rypma, B., Berger, J. S., & D’Esposito, M. (2002). The influence of working-memory demand and subject performance on prefrontal cortical activity. Journal of Cognitive Neuroscience, 14 (5), 721–731. 10.1162/08989290260138627

Silva, M., Baldassano, C., & Fuentemilla, L. (2019). Rapid Memory Reactivation at Movie Event Boundaries Promotes Episodic Encoding. The Journal of Neuroscience, 39 (43), 8538–8548. 10.1523/JNEUROSCI.0360-19.2019

Spreng, R. N., Mar, R. A., & Kim, A. S. N. (2009). The Common Neural Basis of Autobiographical Memory, Prospection, Navigation, Theory of Mind, and the Default Mode: A Quantitative Meta-analysis. Journal of Cognitive Neuroscience, 21 (3), 489–510. 10.1162/jocn.2008.21029

Thomas Yeo, B. T., Krienen, F. M., Sepulcre, J., Sabuncu, M. R., Lashkari, D., Hollinshead, M., Roffman, J. L., Smoller, J. W., Zöllei, L., Polimeni, J. R., Fischl, B., Liu, H., & Buckner, R. L. (2011). The organization of the human cerebral cortex estimated by intrinsic functional connectivity. Journal of Neurophysiology, 106 (3), 1125–1165. 10.1152/jn.00338.2011

Thompson, G. J., Pan, W.-J., Magnuson, M. E., Jaeger, D., & Keilholz, S. D. (2014). Quasi-periodic patterns (QPP): Large-scale dynamics in resting state fMRI that correlate with local infraslow electrical activity. NeuroImage, 84, 1018–1031. 10.1016/j.neuroimage.2013.09.029

Uncapher, M. R., & Wagner, A. D. (2009). Posterior parietal cortex and episodic encoding: Insights from fMRI subsequent memory effects and dual-attention theory. Neurobiology of Learning and Memory, 91 (2), 139–154. 10.1016/j.nlm.2008.10.011

Weigard, A., Angstadt, M., Taxali, A., Heathcote, A., Heitzeg, M. M., & Sripada, C. (2024). Flexible adaptation of task-positive brain networks predicts efficiency of evidence accumulation. Communications Biology, 7 (1), 801. 10.1038/s42003-024-06506-w

Yamashita, A., Rothlein, D., Kucyi, A., Valera, E. M., & Esterman, M. (2021). Brain state-based detection of attentional fluctuations and their modulation. NeuroImage, 236, 118072. 10.1016/j.neuroimage.2021.118072

Zacks, J. M., & Swallow, K. M. (2007). EVENT SEGMENTATION. Current directions in psychological science, 16 (2), 80–84. 10.1111/j.1467-8721.2007.00480.x

